# A FAIR Amplicon Sequencing Workflow for Long-term Environmental Monitoring

**DOI:** 10.64898/2025.12.04.692289

**Authors:** Marie Harmel, Benoit Durieu, Valentina Savaglia, Bjorn Tytgat, Denis Baurain, Annick Wilmotte, Elie Verleyen, Luc Cornet

## Abstract

Antarctica represents one of the last pristine environments on Earth, providing a unique opportunity to to study the effects of climate change and anthropogenic activities. Ice-free areas, such as the inland nunataks of the Sør Rondane Mountains (SRM), host unique terrestrial and lacustrine ecosystems, of which the simplified food webs rely almost exclusively on microbial primary production. Because of their small size, low productivity and hence low biomass, these microbial communities are fragile. Seven SRM sites were selected to be part of the Antarctic Specially Protected Area (ASPA) 179. The MonASPA project has established an environmental monitoring program to evaluate the effectiveness of the management plan approved by the Antarctic Treaty System for these areas. A key component of MonASPA is the long-term monitoring of microbial biodiversity using 16S rRNA amplicon sequencing. To ensure consistent operation over decades, we developed the Reproducible Amplicon Sequencing Pipeline for Antarctic Monitoring (RASPAM), which is built on Apptainer containers and Nextflow workflows. RASPAM implements Amplicon Sequence Variants (ASVs) and Zero-radius Operational Taxonomic Units (ZOTUs) to provide high-resolution taxonomic affiliation. It incorporates, in addition to the SILVA database, a taxonomically curated 16S rRNA database for cyanobacteria and enables comparisons against NCBI databases to facilitate the identification of rare prokaryotic strains in environmental samples. RASPAM is Findable, Accessible, Interoperable, and Reproducible (FAIR) and represents a robust tool for long-term monitoring of microbial communities in Antarctic and other extreme environments.

## Main

Antarctica remains one of the last wildernesses and natural laboratories on our planet, and therefore it offers a unique opportunity to study the effects of climate change and human disturbances on its terrestrial and lacustrine ecosystems. The continent is characterized by some of the most extreme environmental conditions on Earth, namely extremely low temperatures, intense and seasonally variable solar and ultraviolet radiation, and very limited availability of liquid water [1]. The ecosystems in the few ice-free areas, consisting of isolated rocky land such as nunataks and inland mountain chains, host unique and highly endemic biological communities [2]. Among these, the Sør Rondane Mountains (SRM) form a remarkable ensemble of potential ecological refuges [3].

The SRM include a mosaic of terrestrial habitats surrounded by glaciers, along with some rare isolated ponds which, together with cryoconites, represent the only freshwater environments in the region. In these environments, whether terrestrial or lacustrine, food webs are simplified, lacking vertebrates and containing only a few microinvertebrates, and they rely almost exclusively on microbial communities for primary production [4]. The bacterial communities are typically dominated by *Actinomycetota* and *Acidobacteriota* in the terrestrial biofilms on dry morainic substrates, whereas *Cyanobacteriota* prevail in the granitic and marble substrates and in aquatic microbial mats [1].

However, the low productivity and small population sizes of these ecosystems make them particularly vulnerable [5] to anthropogenic pressures such as the introduction of non-native species, trampling, and over-sampling. These disturbances could interact with the effects of global warming and trigger long-lasting changes in the composition of these unique microbial communities. In this context, seven sites within the SRM (**Figure 1**: Perlebandet North and South, Petrellnuten, Tanngarden, Pingvinane, Teltet, and the Yûboku-dani Valley) have been designated within the Antarctic Treaty System as being part of an Antarctic Specially Protected Areas (ASPA) due to their unique environmental, scientific, aesthetic and wilderness values. The MonASPA project aims to assess the effectiveness of the management plan which was developed to protect these values of ASPA 179 through the establishment of a long-term environmental monitoring program based on permanent monitoring plots set up in each of the seven sites (**Figure 1**). Monitoring will include measurements of key environmental parameters (such as soil temperature and humidity, and snow cover), use of passive air samplers, in combination with environmental DNA sequencing, including amplicons of the 16S rRNA gene.

**Figure 1:**
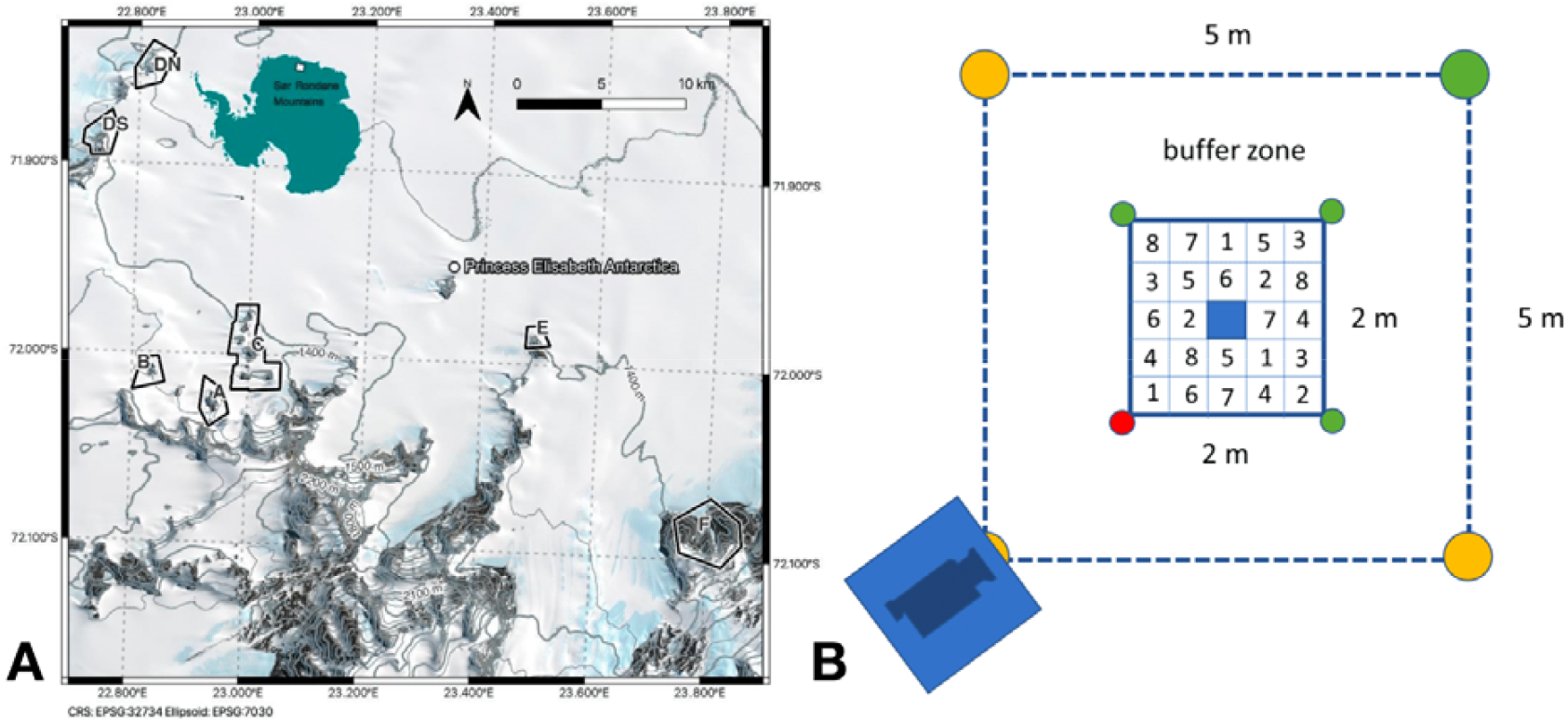
Geographic context of the ASPA sites and the layout of the monitoring plots. **(A)** Map of the Western Sør Rondane Mountains showing the locations of the Princess Elisabeth Station and the seven ASPA sites: Tanngarden Ridge (A), Petrellnuten Nunatak (B), Pingvinane Nunataks range (C), Perlebandet range (DN and DS), Teltet Nunatak (E), and Yûboku-dani Valley (F). The map contains modified Copernicus Sentinel data [2019]; the Antarctica contour is from Wikimedia Commons (https://commons.wikimedia.org/wiki/File:AntarcticaContour.svg) and was released under a CC BY-SA 4.0 license. **(B)** Layout of a single monitoring plot. In each of the seven ASPA sites, three permanent monitoring plots will be installed. Colour-coded stainless steel pegs will mark the plot boundaries to ensure a correct orientation (for sampling), and increase visibility not to disturb the plots. A 1.5 m buffer zone surrounds the central sampling plot to prevent trampling; the entire surface can be accessed from within this zone. The subplot layout (40 × 40 cm) allows researchers to take three samples per campaign in a predefined sequence (eight sampling efforts in total, numbered 1–8, including T0, over 35 years) while avoiding resampling. Taking multiple samples per campaign will compensate for spatial heterogeneity among the communities, and will ensure a representative characterization of their diversity. The central subplot remains unsampled. Additional monitoring devices, such as time-lapse cameras or temperature loggers, can be installed.

One of the main objectives of this genetic monitoring effort is to detect potential introduction or windborne dispersal of non-native microorganisms, and if present, to determine their geographic origin, providing insights into biological connectivity. Given no similar environmental monitoring programs are in place in the Antarctic, the project should build a solid scientific foundation for the long-term assessment of biodiversity and management of Antarctic environments. Yet, since the monitoring will last far beyond the initial phase of the project, the reliability of the results highly depends on the comparability of the data collected, as well as on backward reproducibility of the bioinformatic analyses, to enable meaningful comparisons between datasets sampled at very distant time points. One of the main challenges of the MonASPA project is thus to ensure the bioinformatics reproducibility. Indeed, a survey conducted in 2016 showed that a large majority of researchers (70%) are unable to reproduce the results reported in other studies, and that half of them cannot even reproduce their own analyses [6]. A major cause for this issue lies in the difficulty of maintaining a stable computing environment over time, particularly regarding software libraries and dependencies. Operating-system-level virtualization, through container technologies, offers a solution to preserve a fixed, time-stable, and unbreakable computing environment. Reproducible workflows, which document and record all command lines and parameter settings, further ensure that analyses can be fully reproduced within these controlled environments.

Here, we present an amplicon sequencing workflow designed for long-term environmental monitoring, called the *Reproducible Amplicon Sequencing Pipeline for Antarctic Monitoring* (RASPAM, referring to the ASPA area). Apptainer (formerly known as Singularity) was chosen as the container system for RASPAM due to its compatibility with HPC clusters [7] whereas Nextflow was selected as the workflow management system owing to its built-in version control features [8] and its large, active open-source community. RASPAM follows the same strategy used for the implementation of our GEN-ERA genomic workflows, i.e., the ability to execute every workflow with a single command line, thereby ensuring optimal reproducibility [9]. A detailed user guide is provided to facilitate the use of RASPAM for future monitoring (**Supplementary File1**). Environmental monitoring also requires integration with previous studies, allowing the reuse of earlier collected datasets. Therefore, we tested RASPAM on polar and non-polar datasets generated in 2021, 2023, and 2024, and obtained biological interpretations that are consistent with those reported in the original studies (**Supplementary File 2**).

RASPAM supports state-of-the-art amplicon sequencing methods, with the implementation of alpha (Observed, Shannon, Simpson, Chao1, InvSimpson) and beta diversity metrics (NMDS, MDS/PCoA on Bray–Curtis, Jaccard, Euclidean) (**Figure 2)**. Amplicon Sequence Variants (ASVs) and Zero-radius Operational Taxonomic Units (ZOTUs), that consider each sequence individually, are implemented, providing a high taxonomic resolution in both metrics. This fine taxonomic resolution is essential to determine with confidence whether a taxon has been newly introduced or dispersed between sites during the monitoring period. Indeed, the use of clustered sequences to define Operational Taxonomic Units (OTUs), whether merged at 97% or 99%, does not meet the resolution requirements for meaningful environmental monitoring.

**Figure 2:**
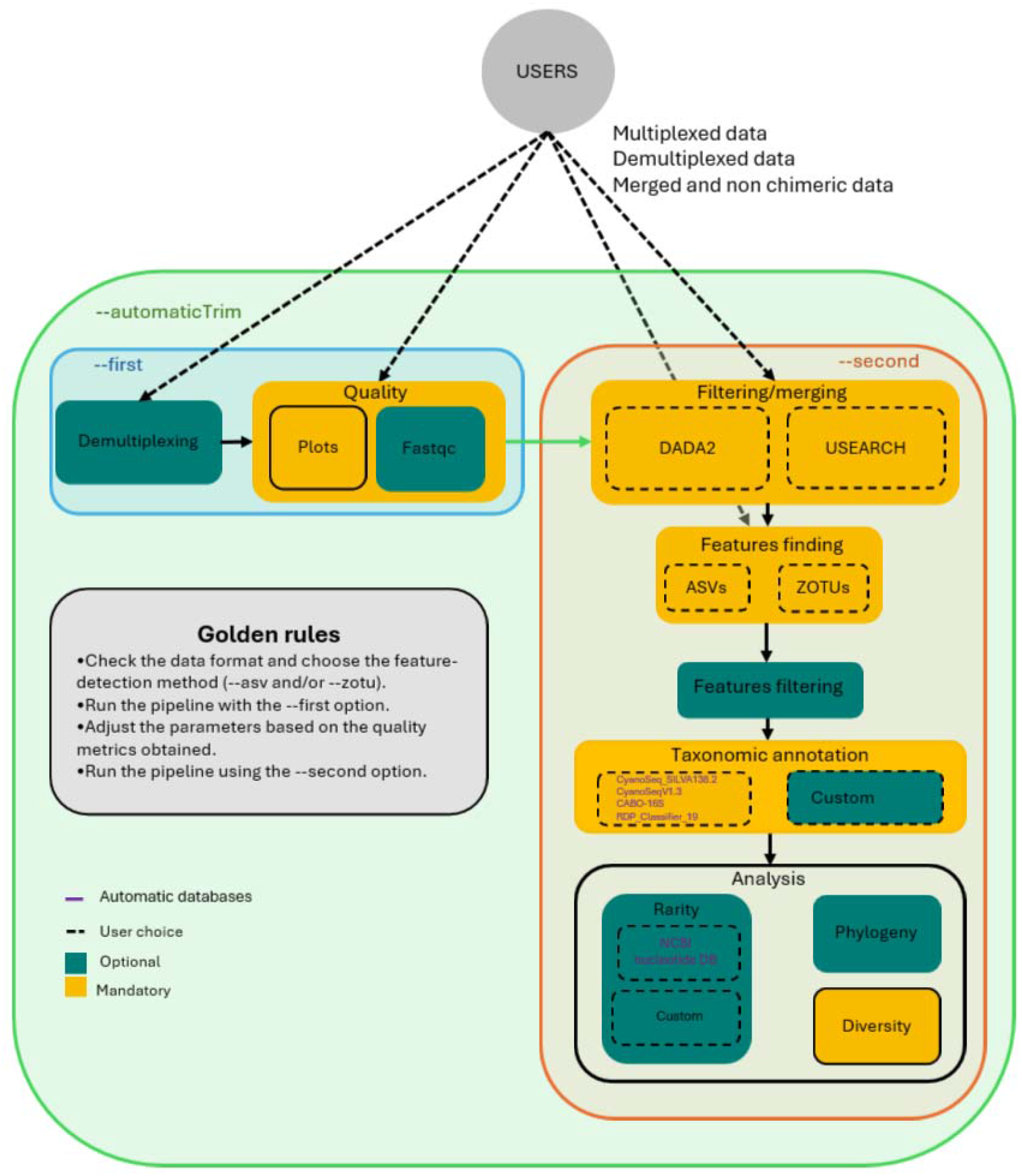
Flowchart of RASPAM.

RASPAM has been developed for prokaryotes using different 16S rRNA databases (**Supplementary file 3**). Cyanobacteria play a major role as keystone taxa in Antarctica, particularly through organic carbon production and nitrogen fixation in microbial mats and biofilms [1]. The diversity of cyanobacteria detected in Antarctica is high, including filamentous orders such as Nostocales, Oscillatoriales, and Pseudanabaenales, as well as unicellular orders such as Chroococcales, Synechococcales, and Gloeobacterales. The current cyanobacterial taxonomy is polyphasic, relying on both genomic and morphological criteria, and is frequently revised. These rapid changes present a greater challenge compared to most other prokaryotes, and can affect the interpretability of data over time. To handle this difficulty, we have incorporated a 16S rRNA database, of complete or nearly complete cyanobacterial sequences, with a taxonomy curated according to the latest Strunecky’s polyphasic classification [10] (**Supplementary file 3)**.

The genomic catalog associated with Antarctica, notably for cyanobacteria, remains limited due to the scarcity of high-quality sequenced genomes, derived from cultures. To address this gap, we have developed a quantitative approach comparing ASV or ZOTU sequences against the NCBI nucleotide databases to detect poorly characterized lineages. RASPAM can determine the rarity and abundance of sequences within samples (**Supplementary file 3)**, facilitating the detection, isolation, and preservation of previously uncharacterized prokaryotic taxa. By applying this approach to samples collected during the MonASPA environmental monitoring program, this will enable the expansion of the Antarctic genomic catalog through the targeted isolation of strains suitable for deposition in public culture collections (e.g., BCCM/ULC; https://bccm.belspo.be/about-ULC).

Antarctica represents one of the last regions on Earth that remain largely unaltered by anthropogenic influence. These ecosystems harbour a unique biodiversity displaying elevated levels of endemism. The protection of the Antarctic environments requires a monitoring procedure that is reproducible, user-friendly, and maintainable over time. We have developed RASPAM for prokaryotes, with the taxonomy particularly optimized for cyanobacteria, by using Apptainer containers and a Nextflow workflow system. RASPAM is Findable, Accessible, Interoperable, and Reproducible (FAIR), and is available as a robust resource that obviously can also be used for monitoring microbial communities in other extreme environments (https://bitbucket.org/phylogeno/raspam/src/main/). However, RASPAM currently is the only amplicon sequencing workflow optimized for cyanobacteria.

## Acknowledgment

This work was supported by a research grant (PDR T.0018.24 OR-OX-PHOT-IN-CYN) from the Belgian National Fund for Scientific Research (F.R.S.-FNRS) to DB. LC is supported by a mandate from the Belgian National Fund for Scientific Research (F.R.S.-FNRS). AW is Senior Research Associate of the FRS-FNRS. The work was also funded by the BELSPO MonASPA project (P4S/25/MonASPA). This is a contribution to the SCAR Research Scientific Programme ‘Integrated Science to Inform Antarctic and Southern Ocean Conservation’ (Ant-ICON). Quinten Vanhellemont (RBINS, Brussels) is thanked for providing Figure 1A. Amandine Bertrand and Julien Ruelle are acknowledged for prototyping an early version of the ZOTU pipeline.

## Contributions

MH and BD coded RASPAM and wrote the Supplementary materials. VS provided part of RASPAM’s preliminary code. MH, BD, VS, DB, AW, EV, and LC wrote the manuscript.

