## Supplementary material for "A FAIR Amplicon Sequencing Workflow for Long-term Environmental Monitoring": Supp file 1

Supplementary file 1: User guide of RASPAM

This supplementary user guide note illustrates how to use the GEN-ERA **RASPAM** tool, from the installation, the job launching command lines, along with a description of each parameter with illustrative output examples.

**Background**

The analysis of microbial communities through amplicon sequencing typically relies on the identification of representative sequences that capture the diversity within a sample. Traditionally, sequences are clustered into Operational Taxonomic Units (OTUs) based on a predefined sequence similarity threshold (e.g., 97%), where each OTU represents a group of closely related sequences assumed to correspond to the same microbial taxon [1]. When a 100% identity threshold is used for clustering, each OTU becomes a ZOTU (Zero-radius Operational Taxonomic Unit), which is conceptually equivalent to an ASV but generated through a different algorithmic framework, specifically a heuristic, abundance-based denoising approach implemented in the UNOISE3 algorithm of the USEARCH program [2].

Alternatively, sequences can be resolved as Amplicon Sequence Variants (ASVs), which represent unique biological sequences inferred after correcting for sequencing errors [3]. ASVs are identified using a probabilistic error model based on the Poisson distribution and an iterative denoising algorithm implemented in DADA2 [3], which separates true biological variation from sequencing noise. This method provides single-nucleotide resolution and allows direct comparison across datasets without the need for arbitrary clustering thresholds [4].

Although ASVs and ZOTUs tend to provide the same type of output, because of the distinct computational principles, they capture different aspects of microbial diversity and can therefore be used alone or in combination.

With these ZOTU and/or ASV sequences identified, filtered by their abundance and/or taxonomies, users can visualize the diversity and also find rare sequences (i.e., not necessarily low-abundance, but lacking close matches in BLAST searches).

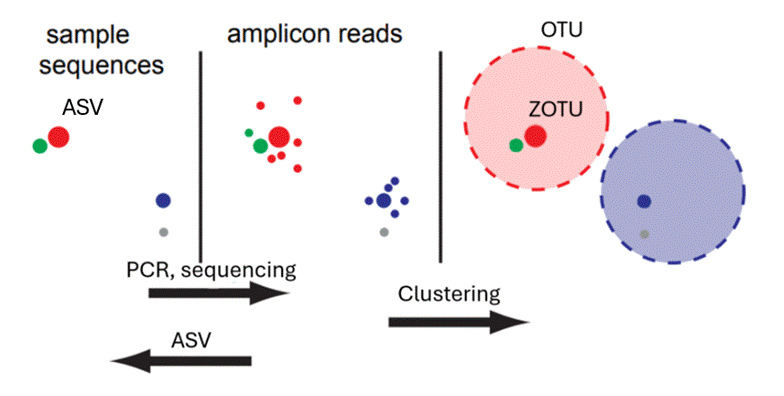

Figure S1: adapted from Callahan et al. (2016)

### **Help**

Mandatory arguments:

--data	DIRECTORY containing the amplicon sequencing data.

one of these arguments:

--first	If enabled, the pipeline will run the first part of the analysis.

--second	If enabled, the pipeline will run the second part of the analysis.

--automaticTrim	If enabled, the pipeline will run entierely with default filtering parameters.

Optional arguments:

quality assessment:

--fastqc	Default NULL. If activated, the pipeline will also run FastQC during the quality plot computing.

--aggregate	Default 'FALSE'. If 'TRUE', display aggregate quality profile plot instead of individual.

--ncol	Default 2. Number of plots' rows displayed on one page of the pdf.

--nrow	Default 3. Number of plots' columns displayed on one page of the pdf.

primers:

--primer_orientation	Default: 'standard'. Specifies the orientation of primers within the paired-end reads. 'standard': the forward primer is present in R1, and the reverse primer is present in R2 (the usual configuration). 'f_r': both the forward and reverse primers are present in each read (R1 and R2).

--forward/--reverse	Default: null/null. String representing the primer sequence that will reorganize the R/F fastq files when in the 'f_r' orientation.

filtering/merging:

asv

--truncQ	Default: 2. The read will be truncated at the first base with a quality score below this value.

--trimLeftF/R	Default: 0/0. Number of nucleotides to remove from the start of each read. F also used for zotu trimming.

--trimRightF/R	Default: 0/0. Number of nucleotides to remove from the end of each read.

--truncLenF/R/_zotu	Default: 0/0. Read truncation lengths for F/R reads, reads shorter than this are DISCARDED. Will run AFTER any of the trimming options.

--in_silico_cutting_min/max	Default 0/10000. Minimum/maximum length of ASV to keep after merging. This is to remove non-specific priming. Set according to your merged amplicon length (length_range_of_ASV.csv in the data/ directory).

--minLenF/R/_zotu	Default: 20/20. Minimum allowed length for F and R reads after trimming and truncation. Reads shorter than this value are discarded.

--matchID	Default: 'TRUE'. Specifies whether to match read IDs during ASV filtering. When you don't have standard header format in your fastq files, set this parameter to 'FALSE'.

zotu

--truncLen_zotu	Default: 200. Minimum allowed length threshold for merged reads (ZOTU workflow).

--forward	Forward is the parameter that is used for reorganize the fastq files but ALSO used to TRIM by its number of character the left side of the merged reads in the ZOTU workflow.

shared

--overlap	Default: 16. Minimum overlap length when merging paired-end reads. Pairs with an overlap shorter than this value will be discarded.

--max_mismatch_merging	Default: 5. Maximum number of mismatches allowed within the overlapping region.

--max_error_F/R	Default: Inf/Inf. Maximum number of expected errors allowed for F and R reads. max_error_R is used for the merged reads (ZOTU workflow).

chimera, denoising, finding feature:

--nderep	Default 1e+06. The maximum number of reads to parse and dereplicate at any one time when learning the error in ASV. This controls the peak memory requirement so that large fastq files are supported.

--learn_number	Default 1e8. The minimum number of total bases to use for error rate learning in ASV.

--pooling_method	Default: 'FALSE'. Specifies how samples are pooled during sequence variant inference in ASV. Possibilities: 'FALSE','TRUE,'pseudo'.

--chimeramethod	Default 'consensus'. The method to use for chimera detection. Possibilities: 'consensus', 'per-sample', 'pooled'.

--zotu_method	Default 'both'. Two algorithms can be used to generate the ZOTU table: 'global' (faster) and 'otutab' (more accurate).

--database_blast	Default local. The path to the 16S database used to blast against features found. If local, the database will be downloaded from NCBI with EDirect.

filtering feature:

--feature_min_occurence	Default 0. Minimum total abundance (read count) required for a feature to be retained.

--feature_max_occurence	Default 'Inf'. Maximum total abundance allowed for a feature to be retained. Features with higher counts are removed.

--feature_min_fraction_on_total	Default 0. Minimum overall relative abundance fraction (across all samples) required to keep a feature. If min 1%: 0.01.

--feature_min_fraction_on_sample	Default 0. Minimum overall relative abundance fraction (in at least one sample) required to keep a feature. If 1% requiered: 0.01.

--feature_min_sample	Default 0. Minimum number of samples in which a feature must appear.

--feature_max_sample	Default 'Inf'. Maximum number of samples in which a feature can appear.

--feature_sample_min_reads	Default 0. Minimum number of reads in at least one sample required to keep a feature.

taxonomic inference:

--databases_taxo	Default local. Path to the 16S taxonomic databases DIRECTORY. If local, the database downloaded will be from CABO-16S, CyanoSeqV1.3 (species), CyanoSeqV1.3_SILVA138.2 and RDPtrainset19.

--filter_taxo	Default 'cyano'. Filters the assigned taxonomy to retain only sequences matching the specified lineage filter: 'cyano' to keep only the *Cyanobacteria* phylum, 'cyano_plas' to include plastids, or 'all' to disable any taxonomic filtering.

--top_taxa	Default 100. Maximum number of taxa or lineages to display in diversity plots; high values may reduce readability due to rare taxa.

--min_bootstrap_assign	Default 50. Minimum bootstrap confidence required for taxonomic assignment.

--phylogeny	Default null. If enabled (--phylogeny), the diversity plot will create the rooted and unrooted tree and fastreee trees too.

--outdir	Default GENERA_AMPLICON. The output directory where the results will be stored.

--cpu	Default 1. The number of CPUs to use.

--help	Display this help message and exit.

### **Arguments**

#### **General options:**

##### **--data**

This parameter specifies the input sequencing data directory for the analysis. The pipeline supports both ASV (Amplicon Sequence Variant) and ZOTU (Zero-radius Operational Taxonomic Unit**)** processing modes, which are both designed for paired-end Illumina sequencing data in FASTQ format.

For ASV and ZOTU, the data can already be demultiplexed or not, and for the ZOTU analysis, the data can also be in a merged file (merged.fq) containing all reads, where each read header includes the sample name.

Forward and reverse reads must share the same identifier in their filenames, followed by the pattern *_R1.fastq for forward reads and *_R2.fastq for reverse reads. If the data are not demultiplexed, a corresponding *_barcodes.fasta file must be provided in the same directory. If 2 barcode files are used (reverse and forward), please name them with *_for_barcodes.fasta and *_rev_barcodes.fasta. Demutliplexing will be done with cutadapt.

The pipeline is designed to also support multiple batches in the same run, but only if the correct structure is kept.

Sample renaming can be performed independently of the barcode names by providing a *.txt file containing the desired names. In cases where two barcode files are used (forward and reverse), samples will be renamed using both names separated by a hyphen. This approach is particularly useful for filtering outputs based on the presence of specific forward and reverse reads (e.g., forward=barcode1 and reverse=barcode1_REV). For instance, a sample renamed as barcode1-barcode2rev will not be retained unless this exact combination is specified in the .txt file. This is advantageous when two barcode names are present and only samples matching paired barcodes (e.g., BC1_BC1_rev, BC2_BC2_rev) are intended to be retained, excluding mixed combinations such as BC1_BC2_rev.

For the diversity plots, you can also add a .idm file (tabulated separated file) where you have your sample_name in the first column and the second one, the new name, without a header. Names not found in it will not be displayed in the plots

Examples:

multiplexed data:

Batch-1_barcodes.fasta Batch-1_R1.fastq Batch-1_R2.fastq Batch-2_barcodes.fasta Batch-2_R1.fastq Batch-2_R2.fastq

Batch-1_rev_barcodes.fasta Batch-1_for_barcodes.fasta Batch-1_R1.fastq Batch-1_R2.fastq

Batch-1_rev_barcodes.fasta Batch-1_for_barcodes.fasta Batch-1_R1.fastq Batch-1_R2.fastq Batch-1.txt Batch-1.idm

demultiplexed data:

HW12_S47_L001_R1.fastq

HW12_S47_L001_R2.fastq

YO_2018_2_R1.fastq

YO_2018_2_R2.fastq

merged data:

merged.fq

###### **File formats**

Barcode file (*.fasta)

When using multiplexed data, a corresponding barcode file must be provided for each batch. Each barcode entry must begin with a header line starting with the character >, followed by the sample identifier, and the next line must contain the corresponding barcode sequence.

>sample_1

ACGAGTGCGT

>sample_2

ACGCTCGACA

Renaming file (*.txt)

A space-separated text file used to rename samples after demultiplexing.

The first column contains the original sample prefix assigned during demultiplexing. If two barcode files are used, the forward and reverse identifiers are combined using a hyphen (e.g., sample_for-sample_rev). The second column specifies the desired new sample name.

sample_1-sample_1_rev control_16s

sample_2-sample_2_rev WT

Merged reads file (merged.fq)

For ZOTU analysis, when using pre-merged data, all reads are combined into a single FASTQ file.
 Each read header must include the sample name, typically indicated by the keyword sample= or an equivalent identifier. _ cannot be used in the name.

merged.fq

@M04880:115:000000000-JDN9V:1:1101:12376:1219;**sample=W1-S41-L001**;

CCTACGGGTGGC…

##### **--first**

Run the initial quality assessment part of the workflow (it should be executed before using --second). It generates the directories GENERA_AMPLICON_quality/ and GENERA_PREPROCESS_DATA**/** within the output folder. The resulting files *Rplots_quality*.pdf, *summary.csv*, and *read_length_distributions.pdf* provide metrics to help determine optimal parameters for read merging and trimming.

GENERA_PREPROCESS_DATA/ directory will temporarily host data produced during demultiplexing or merging. Once the full analysis is complete (after running --second), this directory can be safely deleted.

##### **--second**

Run the second feature finding part of the workflow. This step filters reads, generates ASVs and/or ZOTUs, builds the abundance table, assigns taxonomies, performs diversity analyses, and identifies uncharacterized sequences. It should be executed after using --first.

##### **--automaticTrim**

If --first and --second are not given, or --automaticTrim=’on’, executes the complete pipeline automatically with default filtering parameters, performing the same operations as running --first and --second consecutively.

This fast option is not recommended, as default trimming and merging values may not be optimal.

#### **Quality:**

##### **--fastqc**

Default NULL. If activated (write --fastqc), the pipeline will run FastQC’s Andrews, S. (2010)[5] program during the quality plot computing.

##### **--aggregate**

Default NULL. If activated (write --aggregate), display aggregate quality profile plots instead of individual plots.

##### **--ncol/--nrow**

Default 2/3. Number of rows and columns used to display quality plots on one page of the pdf. If not changed, 2*3 plots will be displayed per page.

#### **Demultiplexing**

##### **--primer_orientation**

Default: 'standard'. Specifies the orientation of primers within the paired-end reads. 'standard': the forward primer is present in R1, and the reverse primer is present in R2 (the usual configuration). 'f_r': both the forward and reverse primers are present in each read (R1 and R2). If --primer_orientation=’f_r’, the pipeline will automatically adjust the reads by switching them in the correct file. We do not use the orient.fwd argument of the filterAndTrim function of DADA2 because it doesn’t allow non-ATGC characters (no N possible).

Example of the f_r orientation, both primers are present in the R1 file ~%:

sample.fastq

@M02713:244:000000000-B2KRR:1:1101:8693:1176 1:N:0:NCCAAT

ACTGTACAGTGGGGAATTTTCCGCAATGGGCGAAAGCCTG…

+

CCCCCGGGGGGFGGGGGGGGGFGGGDGGGGGGGGGGGG…

@M02713:244:000000000-B2KRR:1:1101:11095:1176 1:N:0:NCCAAT

CATAAACGCAAGCCTCAACGCAGCGACGAGCACGAGAGC…

+

CCCCCGGGGGEGGFGGGGGGGGGGGGEECGGGGGGG@FG…

##### **--forward/--reverse**

Default: null/null. String representing the primer sequence that will reorganize the R/F fastq files when in the 'f_r' orientation. They are also used to demultiplex ZOTU and remove them at the end with the barcodes by the fastq_strip_barcode_relabel2.py qiime script [6].

nextflow run main.nf --data=’./data’ --forward='GTGCCAGCNGCCGCGGTAA' --reverse='GGACTACNNGGGTNTCTAAT' --primer_orientation='f_r'

38/b1342e] process > CheckBarcodes (1)	[100%] 1 of 1 ✔

[da/b14c82] process > Demultiplexing (Lot-1)	[100%] 1 of 1 ✔

[-	] process > SortingPrimers	[100%] 1 of 1 ✔

[-	] process > SortingPrimersOtherOrientation [100%] 1 of 1 ✔

[GENERA-INFO]: Barcode files found. Proceeding with demultiplexing.

[GENERA-INFO]: Checking unique barcode names for the entierety of batches

[GENERA-INFO]: Primer orientation is not usual: forward AND reverse are present in R1 AND R2. Sorting reads with f et r primers and switch them

[GENERA-INFO]: Primer forward AND reverse are present in R1 AND R2.

[GENERA-INFO]: Removing primers for data in forward reads

[GENERA-INFO]: Demultiplexing data with cutadapt

#### **Filtering/merging:**

##### **--truncQ**

Default: 2. ASV. The read will be truncated at the first base with a quality score below this value. This option is not recommended by the authors of DADA2 [7, 8] they advise to use max_error_F/R instead (see below).

##### **--trimLeftF/R**

Default: 0/0. ASV/ZOTU. Number of nucleotides to remove from the start of each read. For ZOTU, only use the trimLeftF.

##### **--trimRightF/R**

Default: 0/0. ASV/ZOTU. Number of nucleotides to remove from the end of each read. For ZOTU, only use the trimRightF.

##### **--truncLenF/truncLenR/truncLen_zotu**

Default: 0/0/200. ASV/ZOTU. Read truncation lengths for F/R/merged reads, reads shorter than this are DISCARDED. Will run after any of the trimming options.

##### **--minLenF/minLenR/minLen_zotu**

Default: 20/20/50. ASV/ZOTU. Minimum allowed length for F and R reads after trimming and truncation and for merging ZOTU. Reads shorter than this value are discarded.

Example:

ASV	* low quality base

--trimLeftF/R=4/5	=== primers

R1: [====-------------------------------------------*---*****]

R2: [=====--------------------------------------*--*--*********]

--trimRightF/R=5/7

R1: [---------------------------------------*---*****]

R2: [-----------------------------------*--*--***********]

--truncLenL/R=43/43

R1: [---------------------------------------*---]

R2: [-----------------------------------*--*--****]

merging

R1: [--------------------------------------*---]

|<---- Overlap ----->|

[*-*--*---------------------------------]: R2 rc

merged: [--------------------------------------------------------]

ZOTU

R1: [====-------------------------------------------*---*****]

R2: [=====--------------------------------------*--*--*********]

merging

R1: [====---------------------------------------*---*****]

|<--------- Overlap --------->|

[*********--*--*---------------------=====]: R2 rc

trimLeftF=4

[====--------------------------------------------------------=====]

truncRightF=5

[-----------------------------------------------------------=====]

truncLen_zotu=43

merged: [--------------------------------------------------------]

##### **--matchID**

Default: 'TRUE'. ASV. Specifies whether to match read IDs during filterAndTrim. This parameter should be set to FALSE when the FASTQ files do not use a standard header format.

standard header: @SRR25396444.1 M06682:13:000000000-JCHCR:1:1101:9182:1708 length=292

non standard: @SRR12632873.3 3 length=200

##### **--overlap**

Default: 16. ASV/ZOTU. Minimum overlap length when merging paired-end reads. Pairs with an overlap shorter than this value will be discarded.

##### **--max_mismatch_merging**

Default: 5. ASV/ZOTU. Maximum number of mismatches allowed within the overlapping region. Consider increasing it if you see that the ratio between R1/R2 and merged is not sufficient.

##### **--max_error_F/R**

Default: Inf/Inf. ASV/ZOTU. Maximum number of expected errors allowed for F and R reads after trimming. max_error_R is also used for the merged reads of the ZOTU workflow. EE = sum(10^(-Q/10)), so if you want an overall quality of 20 on reads of 150nt, EE=150*(10^(-20/10))=150*0.01=1.5 expected errors.

##### **--in_silico_cutting_min/max**

Default 0/10000. ASV. Minimum/maximum length of the ASV to keep after merging. This helps to remove non-specific priming. Set according to your merged amplicon length (length_range_of_ASV.csv in the data/ directory).

#### **Chimera, denoising, finding feature:**

##### **--nderep**

Default ‘1e+06’. The maximum number of reads to parse and dereplicate in one time when learning the errors in ASV. This controls the peak memory requirement so that large fastq files are supported.

##### **--learn_number**

Default ‘1e8’. The minimum number of total bases used for error rate learning in ASV.

##### **--pooling_method**

Default: 'FALSE'. ASV. Specifies how samples are pooled during sequence variant inference. Possibilities: 'FALSE', 'TRUE’, 'pseudo'.

FALSE: each sample is processed independently (fastest); TRUE: all samples are fully pooled, increasing sensitivity for low-abundance variants but at a higher computational cost; pseudo: an intermediate approach where pooled information improves error estimation without full pooling, offering balance.

##### **--chimeramethod**

Default 'consensus'. The method to use for chimera detection. Possibilities: 'consensus', 'per-sample', 'pooled'. Consensus: hybrid approach where sequences are classified as chimeric only if it is the case for most samples agreed (recommended); per-sample: chimeras are detected independently in each sample, minimizing false positives but possibly missing rare chimeras; pooled: all samples are analyzed jointly, improving detection of shared chimeras across samples but potentially over-filtering rare true variants.

##### **--zotu_method**

Default 'otutab'. Two algorithms can be used to generate the ZOTU abundance table based on the already inferred ZOTUs: 'global' (faster) and 'otutab' (more accurate) and ‘both’.

global: Search for one hit to a database using the USEARCH algorithm. Alignments are global and this approach is faster, suitable for large datasets, but may slightly underestimate the abundance of highly similar variants (<https://www.drive5.com/usearch/manual/cmd_usearch_global.html>).

otutab: The otutab command generates an OTU table by directly mapping reads to OTUs abundance tables directly by mapping reads to ZOTUs with optimized thresholds. This method is more accurate and recommended when precision in abundance estimation is critical. (<https://www.drive5.com/usearch/manual/cmd_otutab.html>).

##### **--database_blast**

Specifies the 16S rRNA reference database used for BLAST-based taxonomic assignment of the detected features (file, not directory). If set to local, the database will be automatically downloaded from NCBI using EDirect, with the following command: esearch -db nucleotide -query "Cyanobacteria[Organism] AND 16S[Title]" | efetch -format fasta > database_blast.fasta. (same results as Cyanobacteriota). If the download process is interrupted before completion, the temporary database file (GENERA-databases/database_blast) remains incomplete. In such cases, the partial file must be manually deleted before restarting the pipeline; otherwise, the subsequent run will assume that the database already exists and will skip the download step, potentially resulting in corrupted outputs.

As of September 2025, this query retrieved 70,544 sequences and required approximately 30 minutes to complete. Right now, providing an NCBI API key does not accelerate the process.

For a custom database, the input file must be in FASTA format, where each entry begins with a header line starting with >, followed by one or more lines containing the nucleotide sequence.

database_blast.fasta

>PX367225.1 Woronichinia naegeliana CBMC691m 16S ribosomal RNA gene, partial sequence

CTTGGGTTGTAAACCACTTTTATCAGGGAAGAAGCTCTGACGGT

ACCTGATGAATAAGCATCGGCTAACTCCGTGCCAG…

When BLAST is performed for each feature against this database, the best hit will be picked and based on the percentage id, the feature will be either sorted in high percentage and low percentage (uncharacterizable) with the 94% threshold.

#### **Filtering feature:**

##### **--feature_min_occurence**

Default 0. Minimum total abundance (read count) required for a feature to be retained in the abundance table. If no filtering feature option is given, automatically set --min_occ=1 to not crash the process.

##### **--feature_max_occurence**

Default 'Inf'. Maximum total abundance allowed for a feature to be retained. Features with higher counts are removed.

##### **--feature_min_fraction_on_total**

Default 0. Minimum overall relative abundance fraction (across all samples) required to keep a feature. If you want a minimum of 1% --feature_min_fraction_on_total=0.01. Both min_fraction and min_occurrence cannot be used simultaneously.

##### **--feature_min_fraction_on_sample**

Default 0. Minimum overall relative abundance fraction (in at least one sample) required to keep a feature. If 1% required: 0.01.

##### **--feature_min_sample**

Default 0. Minimum number of samples in which a feature must appear.

##### **--feature_max_sample**

Default 'Inf'. Maximum number of samples in which a feature can appear.

##### **--feature_sample_min_reads**

Default 0. Minimum number of reads in at least one sample required to keep a feature.

Example:

| feature/sample | S1 | S2 | S3 | S4 | tot |
| --- | --- | --- | --- | --- | --- |
| F1 | 500 | 400 | 600 | 0 | 1500 |
| F2 | 10 | 0 | 0 | 5 | 15 |
| F3 | 1000 | 1200 | 800 | 1000 | 4000 |
| F4 | 2 | 1 | 0 | 0 | 3 |
| F5 | 50 | 100 | 50 | 0 | 200 |

5718

| min_occurence | max_occurence | min-fraction_tot | min_sample | max_sample | F filtered |
| --- | --- | --- | --- | --- | --- |
| 10 |  |  |  |  | F1, F2, F3, F5 |
|  | 20 |  |  |  | F2, F4 |
|  |  | 0.01 |  |  | F1, F3, F5 |
|  |  |  | 3 |  | F1, F3, F5 |
|  |  |  |  | 3 | F2, F4 |
| 10 | 50 |  | 2 | 4 | F2 |

#### **Taxonomic inference:**

##### **--databases_taxo**

Default local. Path to the 16S taxonomic databases (put your database(s).fasta in this directory and .fa for your species database). If set to local, the database downloaded will be from CABO-16S, CyanoSeqV1.3 (species), CyanoSeqV1.3_SILVA138.2 and RDPtrainset19 [9–12].

database_taxo.fasta

>Phylum;Class;Order;Family;Genus;Taxon(;Species)

Sequence

database_taxo_species.fa

>SeqID Genus species

Sequence

##### **--filter_taxo**

Default 'cyano'. Filters the assigned taxonomy to retain only sequences matching the specified lineage filter: 'cyano' to keep only the Cyanobacteria phylum (without any Chloroplast remaining in the lineage), 'cyano_plas' to include plastids, or 'all' to disable any taxonomic filtering.

##### **--top_taxa**

Default 100. Maximum number of taxa or lineages to display in diversity plots; high values may reduce readability due to rare taxa. Maximum is set to 100, colorblind friendly with a mean_dist of 41.88 for normal, 36.42-32.5 for deuteranopia ,protanopia , tritanopia .

##### **--min_bootstrap_assign**

Default 50. Minimum bootstrap confidence required for taxonomic assignment. Higher values increase confidence but may result in more unclassified sequences, while lower values provide broader coverage but less reliability.

##### **--phylogeny**

Default null. When enabled, two types of phylogenies are generated:

From the diversityplot process: Uses the ape and phangorn libraries, producing both rooted and unrooted trees.
 Alignment-based phylogeny: Built using MAFFT for alignment and FastTree for tree inference.

##### **--outdir**

Default GENERA_AMPLICON. The output directory where the results will be stored. If you are using --first and --second arguments, please keep the same outdir as it sometimes needs data present there.

##### **--cpu**

Default 1. Specifies the number of CPU cores to allocate for the analysis. The pipeline is designed to take advantage of parallel processing, as several modules (e.g., filtering, denoising, BLAST searches) support multi-threading. Increasing this value proportionally reduces computation time, depending on available hardware and system load.

### **Usage**

#### **Prerequisites**

To install and run the RASPAM toolbox, the following software are needed.

●	**Singularity/Apptainer (v1.1.5-1.el8)** [13]

●	**Nextflow (v*21.08.0)***[14]

Other versions may work, but the ones listed below have been tested and validated for stability and compatibility.

The Amplicon.job is a file already configured to run the pipeline on a SLURM cluster.

#### **Access**

The repository is accessible at this address: <https://bitbucket.org/phylogeno/raspam/src/main>, and is publicly accessible. Once cloned and paths configured ([config.sh](http://config.sh/)), the job can be configured, with your data paths and desired arguments.

git clone git@bitbucket.org:phylogeno/raspam.git

**./**[**config.sh**](http://config.sh/)

nano Amplicon.job

#### **Classic command line modes**

Amplicon.job can be run like this:

nextflow run main6.nf --data="./data/16S_data/" --zotu --automaticTrim

However, this option is not yet recommended, as the default parameters are not stringent and the validity of the resulting outputs strongly depends on the chosen filtering settings.

Alternatively, these two commands are privileged, with the --first and --second argument.

nextflow run main6.nf --data='./data/16S_data' --zotu --**first**

#after --first correctly runned, launch the second

nextflow run main6.nf --data='./data/16S_data' --zotu --**second**

nextflow run main6.nf --cpu=40 --data='./data/mutliplexed_16S_data' --outdir="results" --zotu --asv --**second**--forward='GTGCCAGCNGCCGCGGTAA' --reverse='GGACTACNNGGGTNTCTAAT' \

--minLen_zotu=100 --max_error_F=3 --max_mismatch_merging=8 --overlap=80 --feature_min_fraction_occurence_total=0.001 --feature_min_sample=11 \

--filter_taxo="all"--top_taxa=100--feature_min_occurence=0

nextflow run main6.nf --cpu=80 --data='./data/16S_data' --outdir="results"--asv --**second**--forward='GTGCCAGCMGCCGCGGTAA' --reverse='GGACTACNNGGGTATCTAAT' \

--truncQ=2--max_error_F=2--max_error_R=5--truncLenF=220 --truncLenR=160 --trimLeft=19 --trimLeftR=20 --max_mismatch_merging=8 --overlap=80 \

--filter_taxo="cyano"--pooling_method='pseudo'--top_taxa=100--feature_min_occurence=10 -resume

##### **Tips**

The progress of the job can be monitored using the command: **tail -f slurm***.out (exit with Ctrl+C).

Outputs are written progressively to the output directory, eliminating the need to browse the work/ directory or wait for job completion to verify whether each process produced the expected results. If inconsistencies are observed, the job can be terminated using **scancel** <jobid>, parameters adjusted, and the workflow restarted with the **-resume** (one ‘-’) option to preserve previously completed progress.

### **Outputs**

All intermediate and final useful data products are stored in the output/ directory. If you need to inspect additional internal files, you can browse the **work/** directory, using the /id/process_id/ structure shown in the slurm log to navigate into each process-specific subfolder. Inside each task directory, **.command.sh** contains the exact command that was executed, and **.command.log** shows everything printed to standard output.

##### **RShiny**

To use it, run: **‘./rshiny_launch.sh --outdir='the outdir you already provided**'’. This script formats the output directory so you can directly copy it to your local RStudio environment (RShiny library). On your local machine, you can transfer it with: ‘**scp -r cluster:/path/to/raspam/outdir local/path**’. You can open the app.R file and run it, a web browser window will open and provides an interface to navigate all results: data visualization, diversity analyses, plot export, and more.

### **Rerefences**

1.	Stackebrandt E, Goebel BM. Taxonomic Note: A Place for DNA-DNA Reassociation and 16S rRNA Sequence Analysis in the Present Species Definition in Bacteriology. *International Journal of Systematic and Evolutionary Microbiology* 1994;**44**:846–849. https://doi.org/10.1099/00207713-44-4-846

2.	Edgar R. Taxonomy annotation and guide tree errors in 16S rRNA databases. *PeerJ* 2018;**6**:e5030. https://doi.org/10.7717/peerj.5030

3.	Callahan BJ et al. DADA2: High-resolution sample inference from Illumina amplicon data. *Nat Methods* 2016;**13**:581–583. https://doi.org/10.1038/nmeth.3869

4.	Tikhonov M, Leach RW, Wingreen NS. Interpreting 16S metagenomic data without clustering to achieve sub-OTU resolution. *The ISME Journal* 2015;**9**:68–80. https://doi.org/10.1038/ismej.2014.117

5.	FastQC. 2010. 2010.

6.	Caporaso JG et al. QIIME allows analysis of high-throughput community sequencing data. *Nat Methods* 2010;**7**:335–336. https://doi.org/10.1038/nmeth.f.303

7.	Edgar RC et al. UCHIME improves sensitivity and speed of chimera detection. *Bioinformatics* 2011;**27**:2194–2200. https://doi.org/10.1093/bioinformatics/btr381

8.	Callahan B. Determining filtering parameters #232. https://github.com/benjjneb/dada2/issues/232. .
