## Supplementary material for "A FAIR Amplicon Sequencing Workflow for Long-term Environmental Monitoring": Supp file 2

Supplementary File 2: Comparison of RAPSAM Across Three Previously Published Datasets

To demonstrate the reliability of RASPAM, we re-analysed a zOTU dataset (Xiao et al. (2021**)** [1]) and two ASV dataset (Gladkov et al. (2024) [2] and Paulson et al. (2023) [3]) to compare the results with the original studies.

### Xiao et al. (2021): zOTU dataset

Xiao et al. (2021**)** [1] investigated the diversity and community dynamics of bacterioplankton based on 54 sequencing runs collected from the same river during two contrasting hydrological seasons: wet and dry**.** Their analysis revealed how seasonal changes influence bacterial composition, functional profiles, and co-occurrence patterns along an urbanization gradient.

Amplicon sequence data were processed using the RASPAM pipeline with parameters consistent to those employed by Xiao et al. (2021). Primers, barcodes, and short sequences were removed by trimming 19 nucleotides from the 5′ end and 20 nucleotides from the 3′ end of both forward and reverse reads (--trimLeftF=19 --trimLeftR=19 --trimRightF=20 --trimRightR=20), and discarding reads shorter than 100 bp (--minLen_zotu=100).

Paired-end reads were merged using standard USEARCH parameters, requiring a minimum overlap of 16 bp and allowing up to 5 mismatches within the overlapping region (--overlap=16 --max_mismatch_merging=5). Reads with low expected quality were removed based on the maximum number of allowed errors per read, calculated as EE per base=10^(−Q/10)^→10^(−20/10)^=0.01. For a read length of 250 nucleotides, the total expected number of errors per read is: 250×0.01=2.5, leading to a conservative threshold of max_error_F = 3.

Taxonomic assignment was performed using the SILVA database (--databases_taxo='./GENERA-databases/silva/'). For co-occurrence network construction, only ZOTUs representing at least 0.01% of the total abundance across all samples (--feature_min_fraction_on_total=0.0001) and present in more than 20% of samples (--feature_min_sample=11, corresponding to 20% of 54 total samples) were retained.

Because this analysis was not focused specifically on cyanobacterial communities, no taxonomic filtering was applied (--filter_taxo='all'), allowing all detected taxa to be retained for downstream analyses.

The analysis was performed with the following commands:

nextflow run main.nf --cpu=40 --data='./data/Xiao_genomes' --outdir="user_case_results_1" --zotu --first --max_mismatch_merging=5 --overlap=16 –minLen_zotu='100' --databases_taxo='./GENERA-databases/silva/'

nextflow run main.nf --cpu=80 --data='./data/Xiao_genomes' --outdir="user_case_results_1" --zotu --second --max_error_F=3 --feature_min_fraction_on_total=0.0001 --feature_min_sample=11

--filter_taxo="all" --feature_min_occurence=0 --trimLeftF=19 --trimLeftR=19 --trimRightF=20 –trimRightR=20 --databases_taxo='./GENERA-databases/silva/'

Comparison with Xiao et al. (2021) shows that RASPAM, run with only two command lines, reproduces the same alpha diversity, taxonomic composition, and PCoA clustering patterns, demonstrating its efficiency and reliability (**Figure 1**).

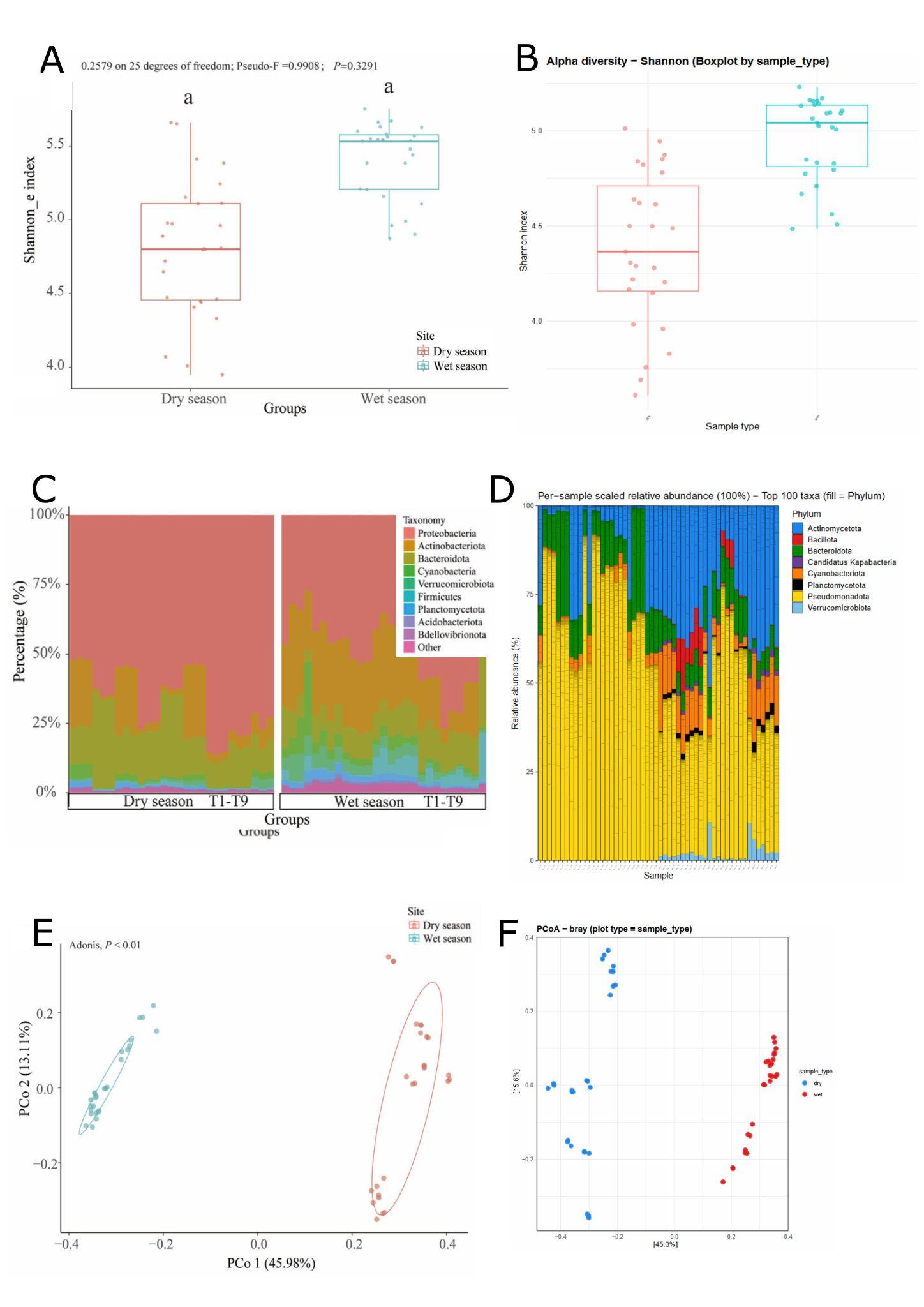

**Figure 1: Comparison between Xiao et al. (2021) and RASPAM analyses. Panels A and B** show boxplots comparing the Shannon indexes between the type of sample (wet and dry) in Xiao et al. (2021) paper and the RASPAM output, respectively. In the dry season, Xiao et al. (2021) reported first and third quartiles around 4.5–5.0, while RASPAM produced slightly lower values (4.2–4.7). A similar pattern is observed for the wet season: Xiao *et al.* reported quartiles around 5.2–5.5, while RASPAM produced values ranging from 4.8–5.1. In both analyses, the median of the wet season is positioned close to the upper quartile, and the upper extreme remains relatively close. **Panels C and D** display the scaled taxonomic β-diversity of the dominant taxa detected across samples. In the original analysis (panel C (Xiao *et al.*, 2021)), several lineage names follow outdated taxonomy[1]; therefore, the following nomenclature adjustments were applied to enable comparison: *ActinobacteriotaàActinomycetota*, *FirmicutesàBacillota*, *CyanobacteriaàCyanobacteriota,* and *ProteobacteriaàPseudomonadota*. After harmonizing taxonomic labels, both datasets show similar phylum-level compositions. Three differences remain, with the presence of Acidobacteriota and Bdellovibrionota in Xiao et al. (2021), and the detection in our analysis of Candidatus Kapabacteria (classified within the Bacteroidota/Chlorobiota group). RASPAM results reveal a clear compositional shift in the middle subset of samples, corresponding to the emergence of Verrucomicrobiota, Candidatus Kapabacteria, Bacillota, and Planctomycetota, when wet-season samples of the dataset begins, which mirrors what is observed in the original study. **Panels E and F** show the PCoA analyses based on Bray–Curtis distances. The color scheme is inverted relative to the original publication (dry = salmon à blue; wet = cyan à red), but this does not affect the interpretation. Because PCoA axes may flip or rotate without changing the underlying ordination structure, both analyses reveal the same overall clustering pattern distinguishing dry-season from wet-season samples. The proportion of variance explained by the first two axes is also highly comparable between the studies: 45.98% and 13.11% for Xiao *et al.* (2021), and 45.3% and 15.6% for the corresponding RASPAM analysis.

### Gladkov et al. (2024): ASV dataset 1

Gladkov et al. (2024) [2] analyzed prokaryotic communities from glaciers located in the Arctic (Mushketova, IGAN), Antarctic (Pimpirev), and Central Caucasus (Skhelda, Garabashi). Each of the 17 cryoconite samples was processed in four independent DNA extraction replicates and sequenced on the Illumina MiSeq platform, targeting the 16S rRNA V4 region (primers 515F/806R; BioProject PRJNA997912). The Garabashi Cracked_glacier samples of the BioProject were not included in the analysis.

Raw data were retrieved using EDirect, renamed according to their sampling location, and processed with RASPAM using parameters consistent with the original study : quality options, silva database, pooling method, and standard values for the ASV merging (--max_mismatch_merging=0 --overlap=12). ASVs were filtered to retain features with at least 10 occurrences between all samples.

The analysis was performed with the following commands:

nextflow run main.nf --cpu=40 --data='./data/Gladkov_genomes' --outdir="user_case_results_2" --asv --first --databases_taxo='./GENERA-databases/silva/'

nextflow run main.nf --cpu=80 --data='./data/Gladkov_genomes' --outdir="user_case_results_2" --asv --second --databases_taxo='./GENERA-databases/silva/'

--truncLenF=220 --truncLenR=160 --trimLeftF=19 --trimLeftR=20 --truncQ=2 --max_error_F=2 --max_error_R=5 --truncQ=2 --max_mismatch_merging=0 --overlap=12 --pooling_method='pseudo' --filter_taxo="all" --top_taxa=62 --feature_min_occurence=10 --feature_min_sample=5

According to Gladkov *et al.* (2024), the core microbiome shared across the Arctic, Antarctic, and Caucasus glaciers is composed of only four ASVs attributed to *Polaromonas* sp., *Rhodoferax* sp., *Cryobacterium* sp., and *Hymenobacter frigidus* [2]. This is also demonstrated by the abundance analyses performed with RASPAM (**Figure 2**).

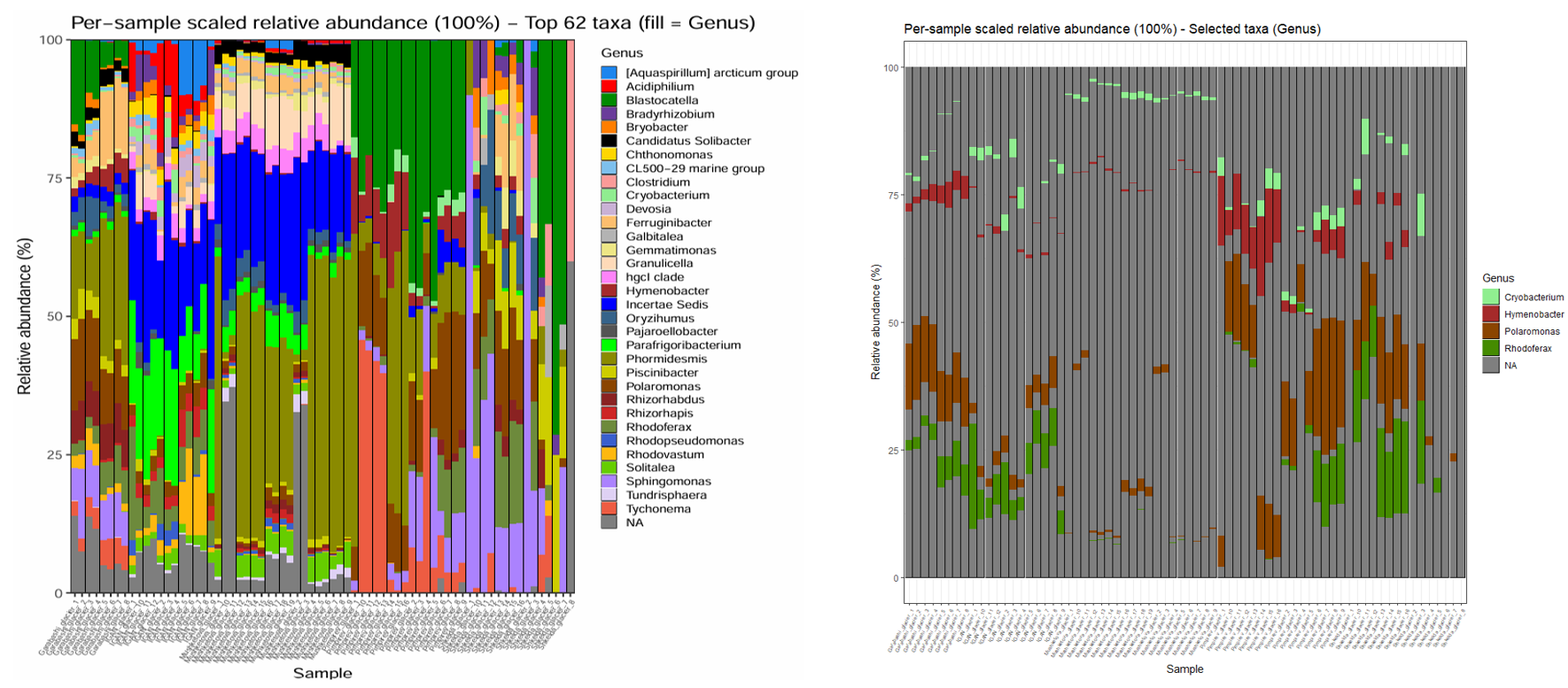
**Figure 2.** **Scaled taxonomic composition (top 100 genera) across all sample**s. The four genera highlighted in the original study: Hymenobacter (dark pink), Rhodoferax (green), Polaromonas (brown), and Cryobacterium (light green), are detected across all glacier sample types. The second panel isolates these genera to better visualize their distribution and relative presence across the different sites.

As noted by Gladkov et al. (2024), the cryoconite microbiomes are composed of several phyla, predominantly Pseudomonadota, Cyanobacteriota, Bacteroidota, Acidobacteriota, and Actinomycetota. Cyanobacteria are particularly dominant in samples from Pimpirev (PAG), PAOr, Garabashi, and Mushketova, whereas they are nearly absent in Skhelda and IGAN, which instead show a higher relative abundance of Pseudomonadota [2]. Using RASPAM, these patterns were automatically reproduced and visualized (**Figure 3**).

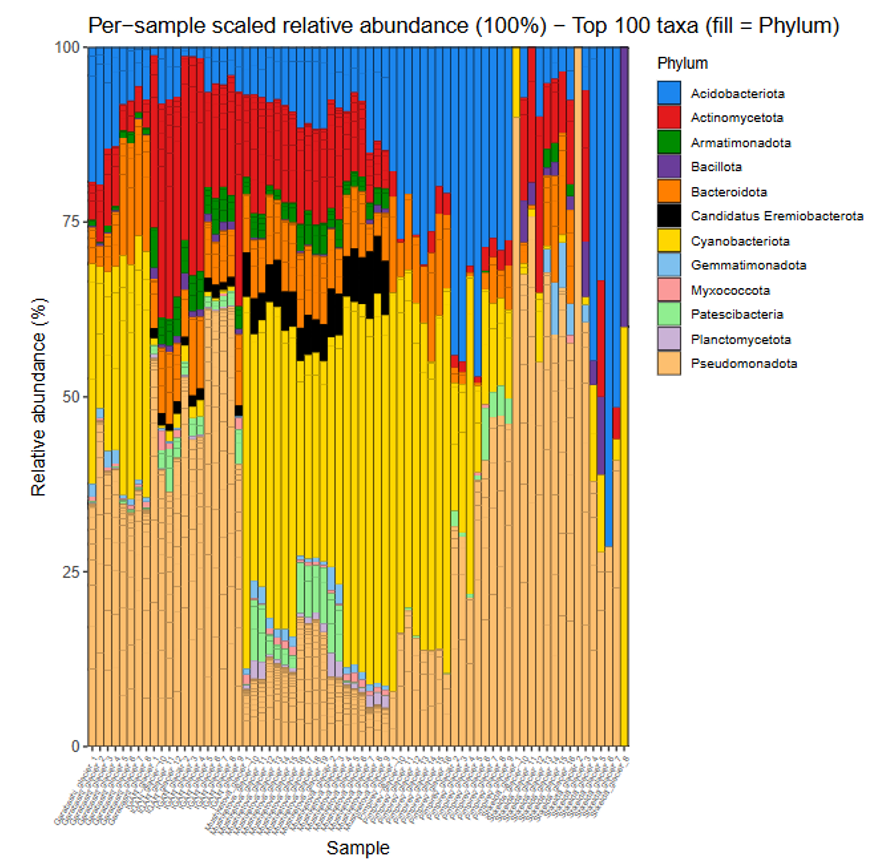

**Figure 3. Scaled taxonomic plot (top 100 phyla) across all cryoconite samples.** The plot highlights the prevalence of Acidobacteriota (blue), Actinomycetota (red), Bacteroidota (orange), Cyanobacteriota (gold), and Pseudomonadota (light orange) across the different glaciers, with distinct lineage structures observable within each site.

The non-metric multidimensional scaling (NMDS) analysis of cryoconite microbiomes in Gladkov et al. (2024) revealed clustering patterns that reflected the geographical proximity of the sampling sites [2]. Beta-diversity analysis plot generated by RASPAM (**Figure 4**) shows that microbial communities from the Arctic glaciers (IGAN, Mushketova) clustered on the left side of the ordination plot, whereas samples from the Antarctic glaciers (PAG, PAOr) grouped distinctly on the right side. The Central Caucasus sites (Garabashi, Skhelda) occupied intermediate positions, bridging the Arctic and Antarctic clusters [2].

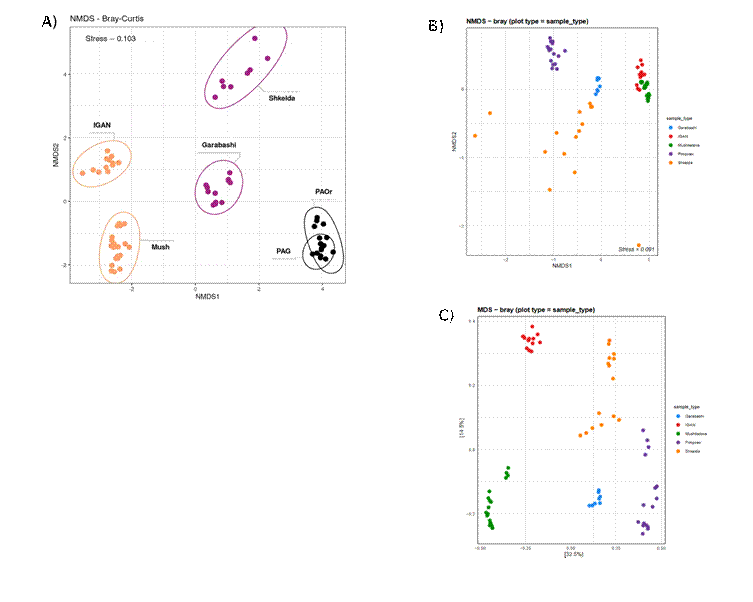

**Figure 4.** **Beta-diversity analysis (NMDS based on Bray–Curtis dissimilarity)**. **Panel A** shows the original ordination from Gladkov et al. (2024), while **panels B** and **C** present NMDS and MDS ordinations generated using RASPAM. Although distinct clustering patterns are visible, no direct quantitative comparison can be made since the data were not CLR-transformed.

### Paulson et al. (2023): ASV dataset 2

A third comparison was performed using the dataset of Paulson et al. (2023) [3], who investigated the microbiome of *Ixodes scapularis* ticks from Canada to assess microbial variation among tissues and to detect potential pathogens [3]. The study analysed four tissue types—salivary glands (sg), gut (g), remaining internal viscera (rem), whole tick (w), and dissected (d) specimens. Although this dataset does not include cyanobacterial taxa, it provides a valuable comparison for testing and illustrating specific components of the RASPAM pipeline, particularly those related to demultiplexing, multiple barcode files, renaming file for diversity plot, handling non-standard header in files.

The raw, multiplexed amplicon data (SRR17194087) were downloaded, and all accompanying metadata were retrieved from the authors’ GitHub repository ([https://raw.githubusercontent.com/damselflywingz/tick_microbiome](https://raw.githubusercontent.com/damselflywingz/tick_microbiome/refs/heads/main/16S_microbiome/data/samples20.txt)). These resources were used to generate the required input files for processing the dataset in RASPAM, including the barcode files, the barcode-to-sample mapping file, and the list of samples flagged as “not kept,” which were excluded accordingly.

Following the authors’ replicate structure (_a, _b, _c), replicate **“**_b**”** libraries were removed, as they were considered problematic in the original study. The remaining replicates _a and _c for each biological sample were retained but not merged *in silico*. Contrary to Paulson et al. (2023) [3], no decontamination filtering (e.g., removal of eukaryotic contaminants or prevalence/frequency-based decontamination) was applied in this processing.

SRR17194087_for_barcodes.fasta

>barcode_10F

CCATCACATAGG

>barcode_11F

GTGGTATGGGAGA

barcode_renaming.txt

barcode_1F-barcode_reverse1 A4_a_16s

barcode_3F-barcode_reverse4 H1_a_16s

barcode_3F-barcode_reverse3 C5_a_16s

sample.idm

A1_a_16s	15_w_a

A1_c_16s	15_w_c

B1_a_16s	31_w_a

The processing parameters were selected to replicate the configuration described in Paulson *et al.* (2023**)** [3] and in the accompanying R scripts available on their [GitHub repository](https://github.com/damselflywingz/tick_microbiome/tree/main/16S_microbiome/reference).

Since ASVs without taxonomic lineage or assigned to Cyanobacteriota were removed in the original analysis, the RASPAM run used the option --filter_taxo=”not_cyano”. Because the FASTQ file headers were non-standard and therefore incompatible with DADA2’s default format, sequence ID matching was disabled with --matchID=FALSE.

Following the filtering criteria reported by Paulson et al.(2023): “Any sample libraries with 7,500 or fewer ASVs were removed. The core bacterial community of *I. scapularis* was defined as the set of ASVs with ≥0.1% relative abundance in at least one library.”, these conditions were translated into RASPAM parameters as:
 --sample_min_reads=7500 and --feature_min_fraction_on_sample=0.001.

For taxonomic assignment, the RDP Train Set 18 database (rdp_train_set_18.fa) was used, and the RefSeq species database was downloaded directly from the authors’ GitHub repository. Because the original diversity plots displayed the 20 most abundant taxa, the visualization parameter was set to --top_taxa=20.

Their primers were specified as ‘forward 515F (5’-GTGCCAGCMGCCGCGGTAA-3’) and reverse 806R (5’-GACTACHVGGGTWTCTAAT-3’)’ [3], which translate in our pipeline as --forward=’ AATGGCGCCGCGACCGTG --reverse=’ATTAGANACCCNNGTAGTC’.

The analysis was performed with the following commands:

nextflow run main.nf --cpu=50 --data='./data/genomes_3' --outdir="user_case_results33" --asv --first --databases_taxo='./GENERA-databases/rdp/' --forward=’ AATGGCGCCGCGACCGTG --reverse=’ATTAGANACCCNNGTAGTC’

nextflow run main.nf --cpu=80 --data='./data/genomes_3' --outdir="user_case_results33" --asv --second --databases_taxo='./GENERA-databases/rdp/' --forward=’ AATGGCGCCGCGACCGTG --reverse=’ATTAGANACCCNNGTAGTC’ \

--truncQ=2 --max_error_F=2 --max_error_R=2 --truncLenF=240 --truncLenR=239 --trimLeftF=20 --trimLeftR=19 --max_mismatch_merging=0 --overlap=12 \

--filter_taxo="not_cyano" --pooling_method="TRUE" --top_taxa=20 --feature_min_occurence=4 --feature_min_fraction_on_sample=0.001 --sample_min_reads=7500 --matchID='FALSE'

RASPAM produced results consistent with those reported by Paulson et al. (2023).
In their study, the authors identified three primary “ASVs of concern” (AoCs) from the core tick microbiome: Borrelia sp. (ASV3), Borrelia miyamotoi (ASV2), and Anaplasma phagocytophilum [3].

Our analyses recovered the same Borrelia-associated ASVs:

"ASV_2","Bacteria","Spirochaetota","Spirochaetia","Spirochaetales","Borreliaceae","Borrelia","miyamotoi"

"ASV_3","Bacteria","Spirochaetota","Spirochaetia","Spirochaetales","Borreliaceae","Borrelia",NA

The third AoC reported in the original paper was not present among the 19 ASVs retained after filtering in our analysis.

We also observed widespread detection of Rickettsia sp. (ASV_1), consistent with the original study [3]. ASV_1 (*Rickettsia buchneri*) was indeed present across all three sampling sites and in both whole-tick and dissected tissue libraries. In our dataset, ASV_1 accounted for 4,424,403 reads, out of a total of 5,170,793 reads across the 19 retained ASVs, representing 85.6% of all sequences (**Figure 5**).

For the taxonomic abundance plots, RASPAM also provided similar results (**Figure 5**), showing the same trend of *Williamsia* sp., *Mycobacterium* sp., *Sphingomonas* sp., *Pseudomonas* (**panel 1 and 2**) and *B. miyamotoi* obvious presence in 33_g and 33_sg samples (**panel 1 and 3**).

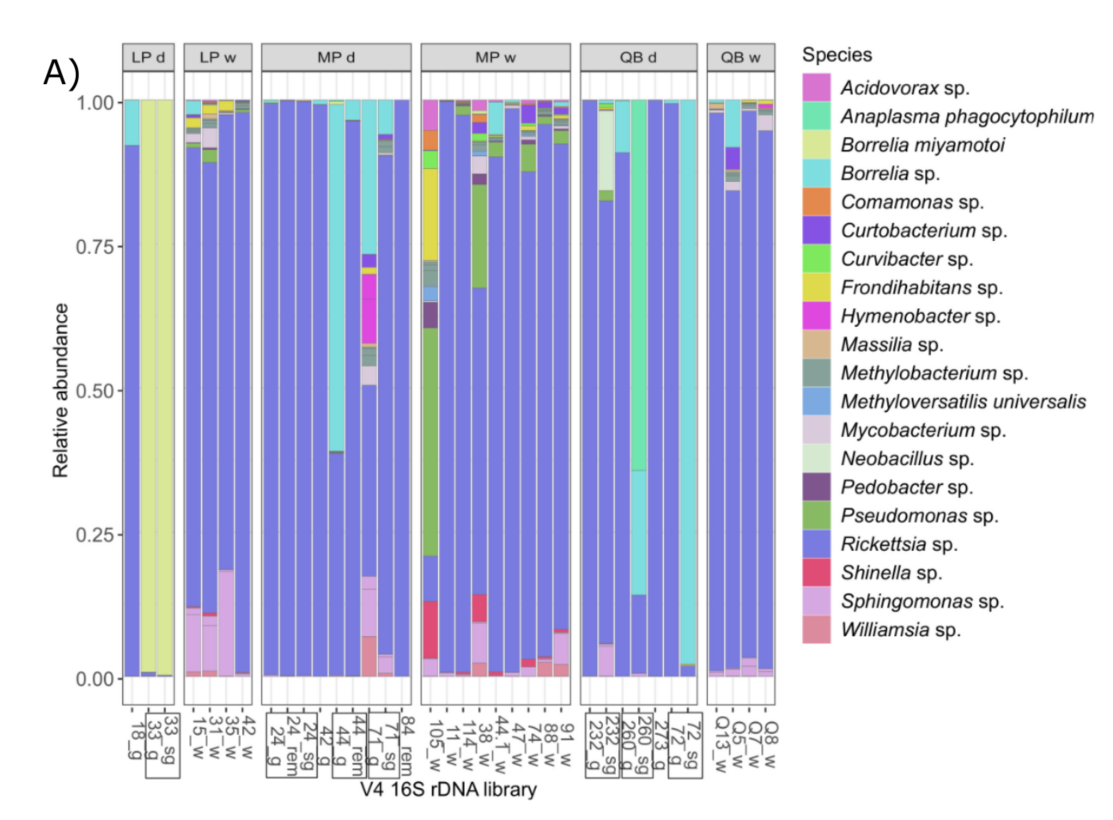

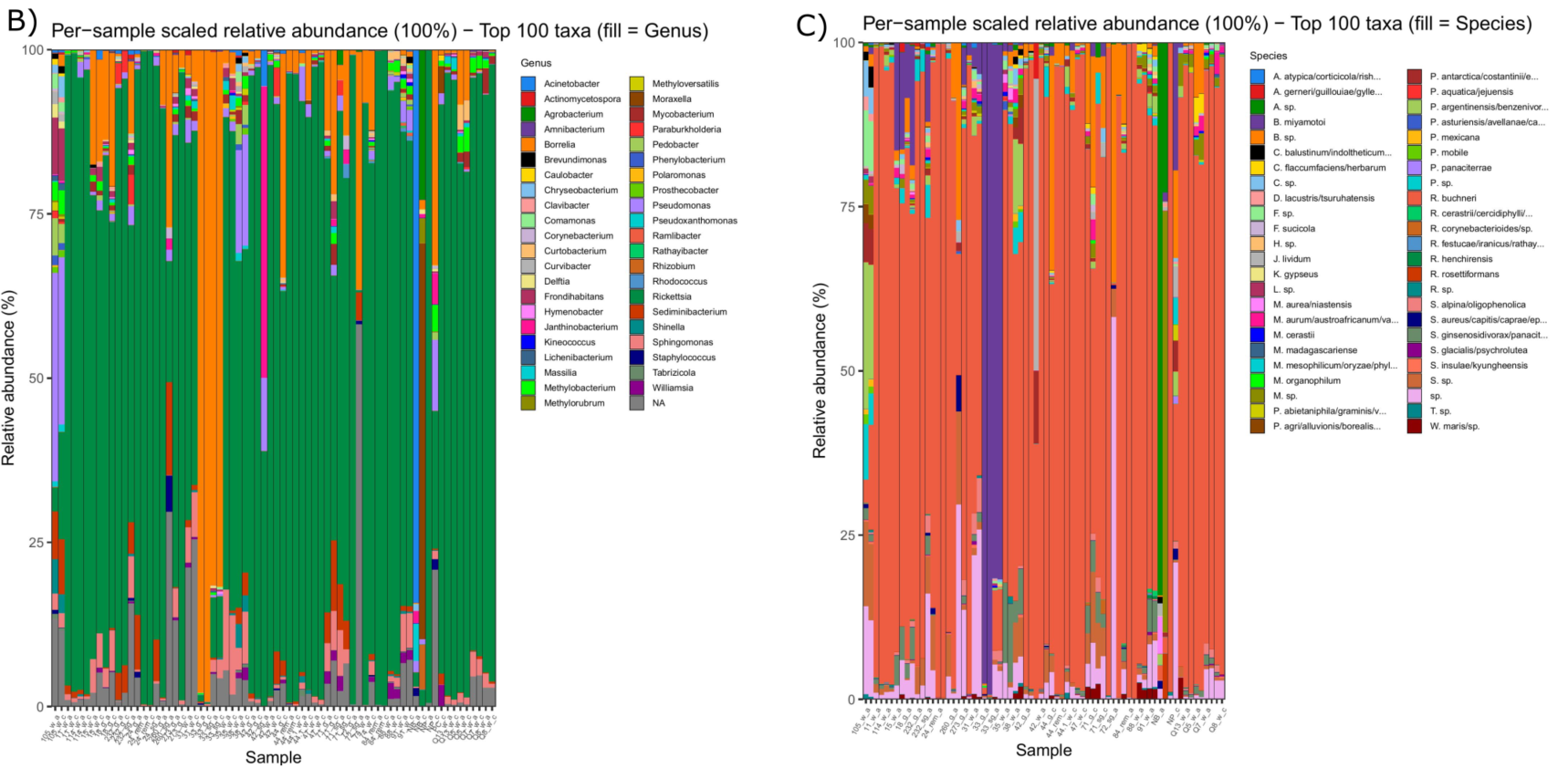

**Figure 5**. **Taxonomic assignments refined to the genus or species level**s. The **panel A** corresponds to the original figure from Paulson et al. (2023), while the two subsequent panels were generated with RASPAM. Each plot displays the 20 most abundant taxa, with relative abundances scaled for visual comparability. In **Panel A**, *B. miyamotoi* is detected at >99% relative abundance in both the 33_sg (salivary glands) and 33_g (midgut) samples. In **Panel B**, several taxa consistently appear in the core microbiome, similar to Panel A: *Williamsia* sp. (purple), *Mycobacterium* sp. (light purple/red-brown), *Sphingomonas* sp. (light purple/light salmon), and *Pseudomonas* sp. (green/light purple), each present in two or more sampling sites [3]. In **Panel C**, *B. miyamotoi* (purple) is widely detected in both the 33_g and 33_sg samples.

With RASPAM, the ASV corresponding to Paulson et al.’s ASV7 is identified as ASV_11, as no ASVs were removed during some decontamination stages.

ASV_11 was successfully detected but, consistent with the findings of Paulson et al.(2023), it could not be taxonomically classified beyond the kingdom *Bacteria* when using the RDP training set and the species-level reference databases integrated in this pipeline[3]:

ASV_11,"Bacteria",NA,NA,NA,NA,NA,NA

CCCATTTGCTACTTTAGCTTTCATCTATCAATGTCAGATATAAACTAGTTTAATACTTTCGTTTATGGCCTTCTTTAATGTATATAAATCATATTTCACCATTATTCAAAAAATTCCTTAAACTTATTTTACTCTCAAGTTTATTAATATTAAATTTAATGTTTAAAAATATTATTTTTAAATTTATAATATAATAGAATAAACCATCTAAAGATGCTTTATGCCCAATAATTGTGAATAACGCTTATAT

Our FeatureAbundanceASV process did not return a BLAST match for this ASV because the local BLAST database contained exclusively cyanobacterial 16S sequences. However, when the ASV_11 sequence was submitted to the NCBI online BLASTN search (<https://blast.ncbi.nlm.nih.gov/Blast.cgi>), we obtained results identical to those reported by Paulson *et al.*:

●	Babesia sp. Dunhuang — E-value: 2e−103

●	Babesia sp. Xinjiang — E-value: 2e−103

●	Babesia orientalis — E-value: 8e−97
