## Supplementary material for "A FAIR Amplicon Sequencing Workflow for Long-term Environmental Monitoring": Supp file 3

Supplementary File 3: Materials and Methods

### **Materials and Software Environment**

The first public release of the RASPAM toolbox was implemented and validated using the software versions listed below. All components were executed within a reproducible containerized environment built with Singularity/Apptainer (*v1.1.5-1.el8*) and managed via Nextflow (*v21.08.0*).

Software and dependencies

●	QIIME (v1.9.1) used for filter_otus_from_otu_table.py and fastq_strip_barcode_relabel2.py [1].

●	Cutadapt (v5.0)[2].

●	USEARCH (v11.0.667_i86linux64) [3].

●	EDirect (v23.9) [4].

●	Bio-MUST-Core (*v0.252040*)

●	Perl (*v5.43.2*)

●	FastTree (*v2.1.11 SSE3*) [5]

●	MAFFT (*v7.505*) [6]

●	BLAST+ (*v2.12.0*) [7]

**Methods**

The methods described below represent the complete RASPAM workflow. Depending on the input data and user-specified arguments, some modules may be automatically skipped to adapt to the dataset structure and analysis mode.

#### **Amplicon Sequence Variant (ASV) unique steps of the workflow**

**Demultiplexing**

When primer sequences are provided and correspond to the *.barcodes.fasta format, RASPAM performs demultiplexing using Cutadapt (v5.0)[2]. One or two barcode (one for forward, one for reverse) files can be specified and the following parameters are applied: -m 50 --error-rate 0 --trimmed-only --times 5. If a sample renaming file (.txt) is provided, sample identifiers are automatically updated to match the new naming scheme. Those that do not match are discarded.

**Primers handling**.

In the standard configuration (the forward primer in R1 and the reverse primer in R2), primer trimming is performed by Cutadapt [2] where the forward and reverse primers are provided respectively via -g and -G: --trimmed-only --no-trim -m 50 --error-rate ${error_rate} --action=retain. When primers are present in both reads (primer_orientation set to 'f_r'), the pipeline performs 2 sorting processes: the first sorts like above and the second, with inverted primer positions (-g reverse, -G forward). At this stage, it switches the reads to restore the correct pairing between R1 and R2.

**Quality filtering**

ASV read filtering is performed with the filterAndTrim() function from the DADA2 package (1.34.0) [8], using the following customizable parameters: truncQ, truncLenF / truncLenR, trimLeftF / trimLeftR, trimRightF / trimRightR, minLenF / minLenR, max_error_F / max_error_R. When --primer_orientation='f_r', the pipeline automatically uses the --orient argument to prevent mismatches between forward and reverse reads. If FASTQ headers are non-standard, the option --matchID can be used to disable read ID matching during filtering.

**Error Learning, Merging, and Chimera Removal**

Error rate learning is performed prior to merging, using a dereplication step with a customizable number of reads (--nderep). Error models are trained using the DADA2 dada() function[8], with pooling behavior controlled by the --pooling_method parameter (FALSE, TRUE, or pseudo). Paired-end reads are merged using the user-defined --overlap and --max_mismatch_merging values. After merging, sequences are optionally filtered in silico based on their length to remove amplicons that fall outside the expected length range for the targeted 16S region. Chimera detection and removal are carried out with the removeBimeraDenovo() function [8], according to the selected method parameter (consensus, per-sample, or pooled).

#### **Zero-radius Operational Taxonomic Unit (ZOTU) unique steps of the workflow**

**Demultiplexing and Read Merging**

For the ZOTU workflow, demultiplexing is performed after paired-end reads are merged with using QIIME’s fastq_strip_barcode_relabel2.py script [1]. Read merging itself is carried out with the usearch11 -fastq_mergepairs () function [3], using the parameters --max_mismatch_merging, --overlap, and --minLen_zotu. They are merged only if they share at least 90% identity across the overlapping region.

**Primers**

Once merging is complete, if the dataset is multiplexed, barcodes and primers are removed using the QIIME utility fastq_strip_barcode_relabel2.py [1] twice (one with the forward, one with reverse primer) [1]. This script also renames sequence headers to keep track of the sample in the merged reads. If the data were already demultiplexed, this step is skipped because the merging step already renamed the header with samples, and barcodes are not present.

**Filtering**

For demultiplexed datasets, filtering uses the same principles as the ASV workflow: reads are processed according to the parameters: truncLen_zotu, trimLeftF / trimRightF, --max_error_R with the usearch11 -fastq_filter function [3]. For multiplexed datasets, both output sets from the demultiplexing step are filtered separately and do not use trimLeftF, as primers were already removed. The reverse reads are reverse-complemented before concatenation with the forward reads.

**Singleton Removal, Denoising, and ZOTU Table Generation**

After filtering, singleton sequences (features with fewer than 2 occurrences) are discarded , denoising is then performed with the UNOISE3 algorithm[9] implemented in USEARCH[3], producing zero-radius OTUs (100% identity clusters). Abundance tables are generated using one of two algorithms of usearch11, depending on the zotu_method: global (faster, aligns reads to existing ZOTUs using the usearch_global algorithm[10]), otutab (more accurate, directly maps reads to ZOTUs to construct abundance tables <https://drive5.com/usearch/manual/cmd_otutab.html>).

#### **Common steps of the workflow**

**Quality assessment**

Quality assessment is performed both before and after filtering for ASV and ZOTU feature-finding workflows. The profileQualityPlot() function of DADA2 [8] generates per-sample or aggregate read quality profiles for both R1 and R2 reads. Users can customize the display parameters (--ncol, --nrow) and optionally enable FastQC for additional quality summaries. When the --aggregate option is set to TRUE, a single combined quality plot per read direction is produced instead of individual plots per sample.

**Filtering the feature table**

After sequence inference, the resulting feature abundance table is filtered to remove low-quality or non-informative features. Filtering uses the following parameters: --feature_min_occurrence, --feature_max_occurrence, --feature_min_fraction_on_total, --feature_min_sample, --feature_max_sample. These thresholds control minimum abundance, prevalence across samples, and total relative frequency. Filtering is performed using the QIIME script [1]filter_otus_from_otu_table.py, which operates on a BIOM-format table to retain only relevant features for downstream analysis.

**Taxonomy, Diversity, and Phylogeny**

Taxonomic assignment is performed using DADA2’s assignTaxonomy() function ([8]), based on user-specified or automatically downloaded reference databases (e.g., SILVA (v138.2) [11], CABO-16S [12], CyanoSeq (v1.3)[13], RDPClassifier (v19) [14], or custom databases). The minimum bootstrap confidence can be set with --min_bootstrap_assign (default: 50) and the reverse complement is also checked.

wget https://zenodo.org/records/13910424/files/CyanoSeqV1.3_SILVA138.2_dada2.fastq.gz

wget -O CyanoSeqV1.3.xlsx "https://zenodo.org/records/13910424/files/CyanoSeqV1.3.xlsx"

wget -O CABO-16S.fasta.gz "https://ndownloader.figshare.com/files/49952913"

After taxonomic assignment, results are optionally filtered by lineage using the --filter_taxo parameter: 'cyano', 'cyano-plas', 'all', ‘not_cyano’ which filter respectively only Cyanobacteria, Cyanobacteria and plastids , disable lineage filtering or not cyanobacteria.

Diversity analyses include α-diversity metrics: Observed richness, Shannon, Simpson, InvShannon and Chao1 indices. β-diversity metrics: Ordination (NMDS, MDS, PCoA) and distance-based clustering (bray, jaccard, euclidean), with “square-root relative abundance transformation" possible.

Relative abundance plots for the top N taxa (--top_taxa customizable) and rarefaction curves are also calculated. Phylogenetic inference is conducted in two possible ways: one with R package (DECIPHER (2.22.0) [15], pharagorn (2.12.1) [16], ape (5.8.1)[17] , ggplot2 (4.0.0) [18]) with rooted and unrooted tree, or by the aligning sequences with MAFFT [6], constructing a maximum-likelihood tree using FastTree [5], and refining the output with format-tree.pl (Bio-MUST-Core utilities REF) to map features to their respective taxonomic lineages.

**BLAST Annotation**

To detect uncharacterized or low-similarity features, RASPAM performs BLAST [7]searches against a custom 16S rRNA reference database or an automatically downloaded database from NCBI. Each sequence is compared using blastn ( blastn -query ${chunk} -db ${dbprefix} -num_threads ${task.cpus} -outfmt "6 qseqid sseqid pident length mismatch gapopen qstart qend sstart send evalue bitscore" -out blast_${file_type}__${chunk.baseName}.tsv) and the best hit is retained, its percentage identity, name of the match and lineage of the taxonomic databases. A threshold of 94% sequence similarity identifies potentially novel or unclassified taxa.
